# Computational fluid dynamics simulation of changes in the morphology and airflow dynamics of the upper airways in OSAHS patients after treatment with oral appliances

**DOI:** 10.1101/688879

**Authors:** Song Baolong, Li Yibo, Sun Jianwei, Qi Yizhe, Li Peng, Li Yongming, Gu Zexu

## Abstract

**Objectives:** To explore the changes of morphology and internal airflow in upper airways (UA) after the use of oral appliances (OAs) in patients with obstructive sleep apnea hypopnea syndrome (OSAHS), and investigate the mechanisms by which OAs function as a therapy for OSAHS.

**Methods:** Eight OSAHS patients (all male, aged 37-58, mean age 46.25) underwent CT scans before and after OA use. Then, computational fluid dynamics(CFD) models were built on the base of the CT scans using Mimics and ANSYS ICEM CFD software. The internal airflow of the upper airways was simulated using ANSYS-FLUENT and the results were analyzed using ANSYS-CFD-Post. The data were analyzed to identify the most important changes of biomechanical properties between patients with and without OA intervention. Upper airway morphology and the internal airflow changes were compared using *t*-tests and Spearman correlation coefficient analysis.

**Results:** The narrowest area of upper airways was found to be located in the lower bound of velopharynx, where the volume and pressure were statistically significantly increased (P<0.05) and the air velocity was statistically significantly decreased (P<0.05) in the presence of the OA(P<0.05). After wearing OA, pharyngeal resistance was significantly decreased (P<0.05), from 290.63 to 186.25Pa/L, and the airflow resistance of the pharynx has reduced by 35.9%.

**Conclusion:** The enlargement of the upper airway after wearing the OA changed its airflow dynamics, which decreased the negative pressure and resistance in narrow areas of the upper airways. Thus, the collapsibility of the upper airways was reduced and patency was sustained.

## Introduction

The obstructive sleep apnea-hypopnea syndrome (OSAHS) is a common respiratory sleep disorder that is characterized by repeated partial or complete obstruction of upper airway at the end-expiratory phase during sleep^[1]^. OSAHS is a highly prevalent disorder which can have serious effects on daily functioning, social life and general health and may even have potentially life-threatening consequences. Prevalence studies estimate that 24% of men and 9% of women in middle age are affected by OSAHS^[2]^. Furthermore, the prevalence of OSAHS increases with age and in older persons (≥ 65 years) there is a 2- to 3-fold higher prevalence compared with those in middle age (30–64 years)^[3]^. There is increasing evidence that untreated OSAHS is associated with ischemic heart disease, arterial hypertension stroke^[4,5]^, hypertension^[6]^, daytime sleepiness, and road traffic accidents^[7]^. Thus, it is now recognized that OSAHS is a serious public health problem.

Treatment modalities for OSAHS includes non-surgical treatments such as weight loss, nasal continuous positive airway pressure (CPAP) and oral appliance (OA)^[8–10]^, as well as surgical treatments such as uvulopalatopharyngoplasty (UPPP), hyoid myotomy and suspension, mandibular osteotomy with genioglossus advancement, adenotonsillectomy and maxillomandibular surgery^[11–12]^.Since Sullivan et al. first reported the use of CPAP therapy in OSAHS in1981^[9]^, CPAP has been considered as an effective method for the treatment of OSAHS. However, its clinical effectiveness is limited by poor acceptance and tolerance, as well as suboptimal compliance of patients^[13–14]^. Disadvantages of UPPP surgery include the pain and expense of the procedure, but also a relatively poor long-term success rate^[15]^. Since Robin et al. first introduced an intraoral appliance to treat upper airway obstruction in 1934^[10]^, OAs have increasingly been used in the treatment of OSAHS as a viable alternative to CPAP. While the curative effect of OAs was affirmed by increasing numbers of studies^[16]^, the recent research on the specific mechanism by which OAs affect OSAHS is mainly limited to morphological descriptions^[17–19]^. It is generally believed that after wearing an OA, the tissue surrounding the upper airways of OSAHS patients is pulled out, and the upper airways are expanded, so that local stenosis is released or entirely abrogated. Hence, the patient can breathe easily and have a higher blood oxygen saturation, a lower snoring sound or even no snoring. However, what happens to the flow dynamics of the upper airways after wearing the OA is still not clear. This study therefore probed the flow dynamics of the upper airways of patients with OSAHS in order to understand the relationship between the morphology of upper airways and their function. Ultimately, the data gathered here improves the understanding of the pathogenesis of OSAHS and the therapeutic mechanism of OA.

## Materials and Methods

### Participants

This study protocol was approved by the Ethics Review Committee at the College of Stomatology, Fourth Military Medical University. It conforms to relevant national and international guidelines. Written informed consent was obtained from each patient. In this study 8 Chinese adult OSAHS patients, aged 37 to 58 years (mean age 46.25 years), were selected. All OSAHS patients had an AHI index higher than 5 and lower than 40 per hour on standard polysomnography (PSG). Those who had active temporo-mandibular disfunction were excluded from the study as well as patients suffering from untreated caries and periodontal disease, completely edentulous patients and those having an insufficient number of remaining teeth.

### Three-dimensional reconstruction of upper airways

All patients underwent a PSG detection and a CT scan before and three months after wearing the OAs. CT scans from the lower rim of the epiglottis to the supraorbital margin were performed with a spatial resolution of 512×512 pixels and zero space, 1.25mm thickness, to obtain DICOM (Digital Imaging and Communication in Medicine) format images. Scanning was performed with the patients awake, in supine and neutral position during one breath hold at the end of a normal inspiration. All patients were scanned by the same radiology technicians in the Department of Radiology, Fourth Military Medical University, Xi’an, China. The CT scans were used to construct three-dimensional models for the analysis of fluid dynamics.

The DICOM image data of the patients was then loaded into Mimics 10.0 (Materialise NV, Leuven, Belgium). Subsequently, a segmentation of the upper airways was done based on the Hounsfield unit (HU), a measure of the electron density of the tissue, assigned to each pixel in the series of DICOM images. The HU had values from 1024 to 3071. For the upper airways, a range of HU between −1024 to −400 is regarded as very good results^[20]^. The segmented region was successfully converted to a 3D model, smoothed in Magics9.9 (Materialise, Leuven, Belgium), and finally exported for further analysis (Fig 1).

**Fig 1.**
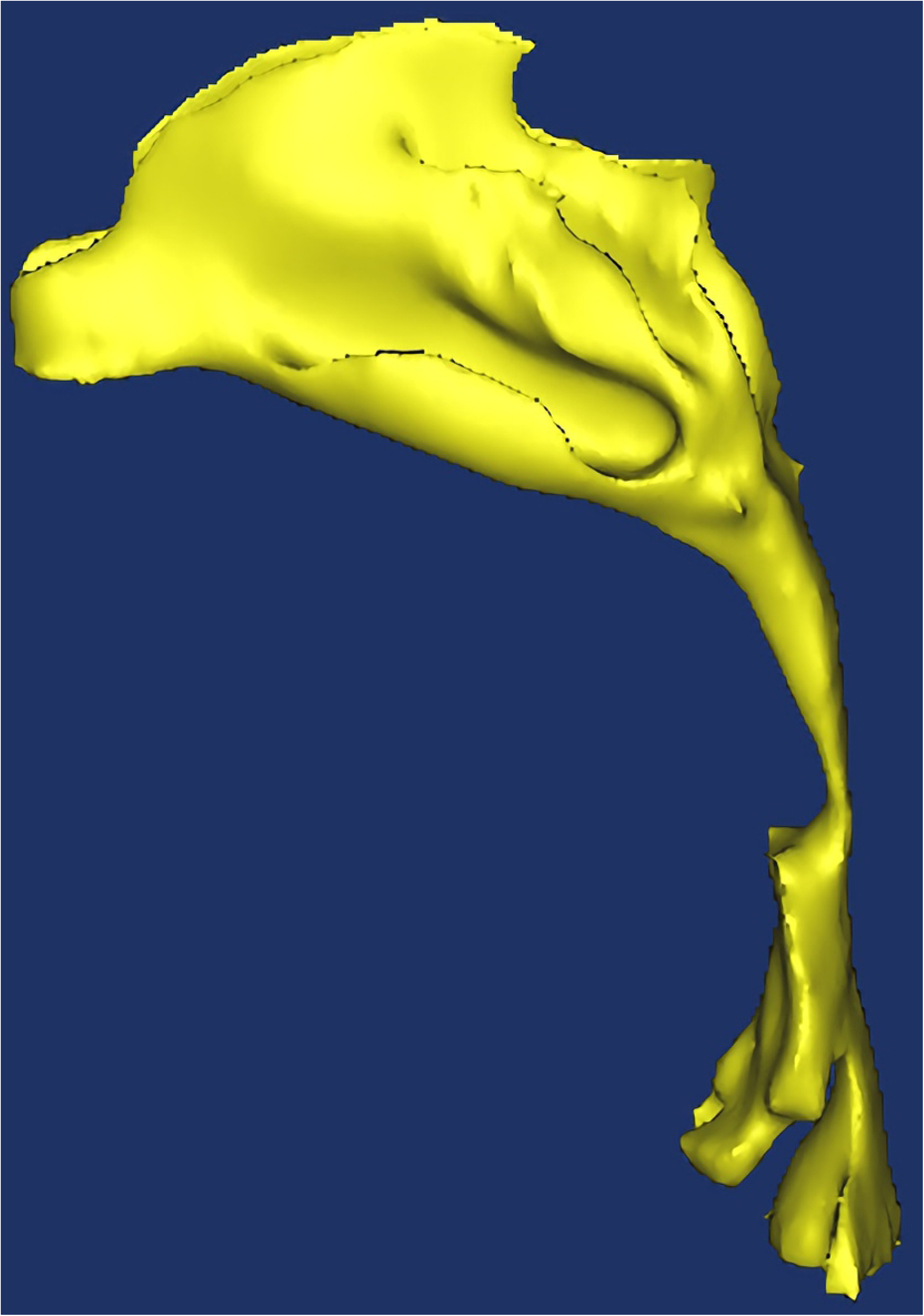
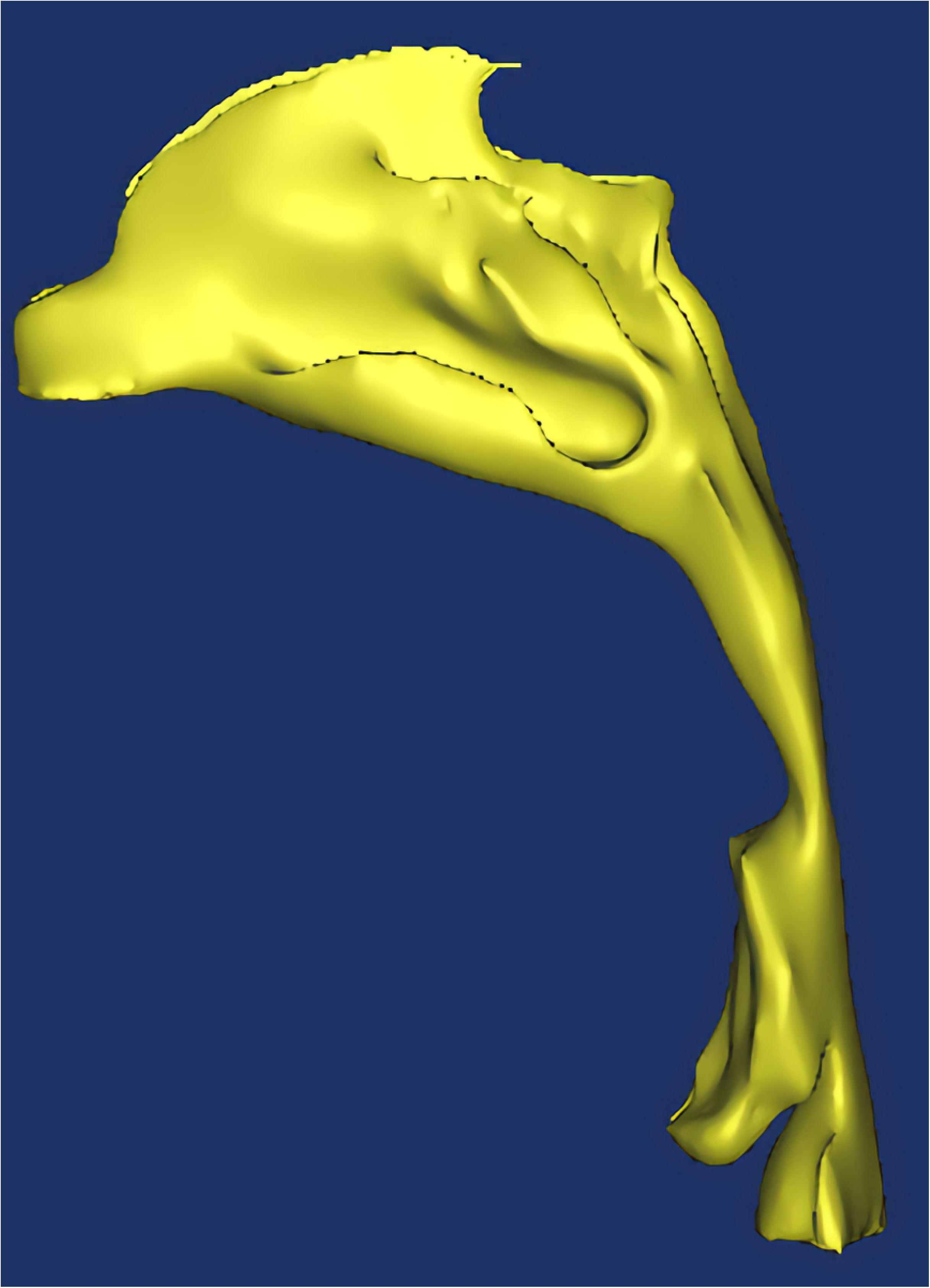
Three-dimensional reconstruction of a patient’s upper airways, without(a) and with (b) an oral appliance.

### Three-dimensional mesh model building

To prepare the segmented upper airway model for CFD calculations, a computational grid was created using ICEM-CFD 14.0 (ANSYS, Canonsburg, PA, USA), in which both global and local mesh controls were used to improve mesh quality (Fig 2), followed by a mesh-independent test.

**Fig 2.**
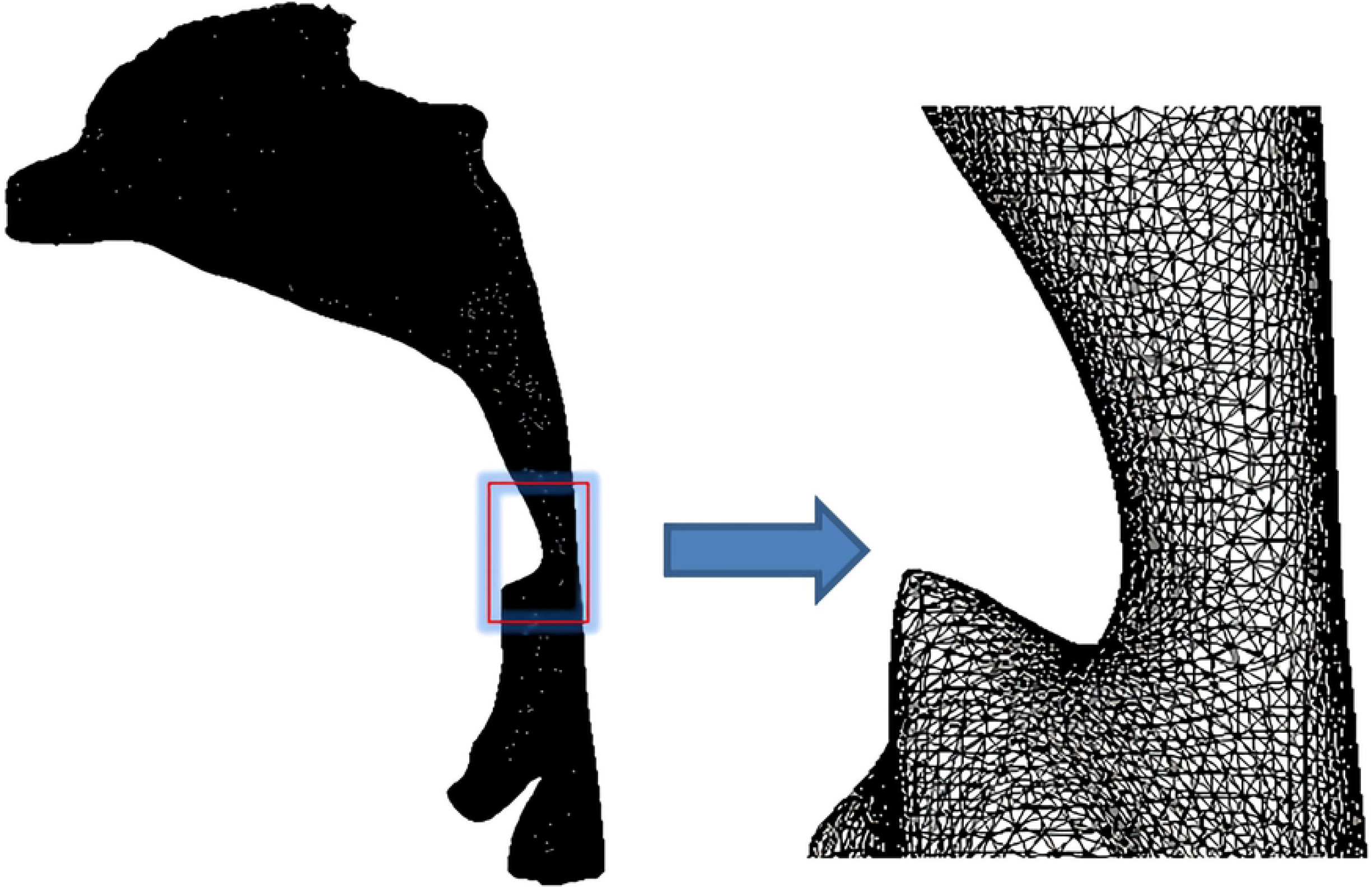
Three-dimensional mesh model of the upper airways.

### Numerical simulation of the upper airways

The ANSYS 14.0-Fluid Dynamics-FLUENT 14.0 commercial code was used to solve the Reynolds averaged Navier–Stokes equations for the steady airflow simulation. The flow was assumed to be steady, homogeneous, incompressible, adiabatic and Newtonian. The density of air was set at 1.225 kg m^−3^, and the viscosity was assumed to be 1.789 x10^-5^ kg m^−1^ s^−1^. We applied the Reynolds averaged Navier-Stokes equation (RNG k-epsilon turbulence model) to simulate the airflow status. A second-order pressure discretization scheme was used for the pressure calculations, and a second-order upwind scheme was used for the momentum and turbulence transport equations. We used the SIMPLE algorithm to solve the pressure–velocity coupling. Atmospheric pressure was imposed at both nostrils, and a constant flow rate of 200 ml/s was applied at the planes in the pharynx. The upper airway wall was assumed to be no-slip (u=v=0). To obtain a better initialization field and convergence acceleration, multigrid initialization was used to optimize the flow field.

The results obtained using Fluent were post-processed using CFD-Post (ANSYS, Canonsburg, PA, USA) to obtain the contours and vectors, and the function calculator was used to acquire values for the pressure gradient, pressure and velocity in cross-sectional planes at specified points in the upper airways.

### Cross-sections selection of the CFD model

The magnitude of airflow velocity in the upper airways was calculated at the five different cross-sections shown in Fig 3. Section 1 constitutes the beginning of the nasopharynx, section 2 is located at the lower bound of the nasopharynx (upper bound of the palatopharynx), and section 3 at the lower bound of the palatopharynx (upper bound of the glossopharynx). Sections 4 and 5 are located the top and the base of the epiglottis, respectively.

**Fig 3.**
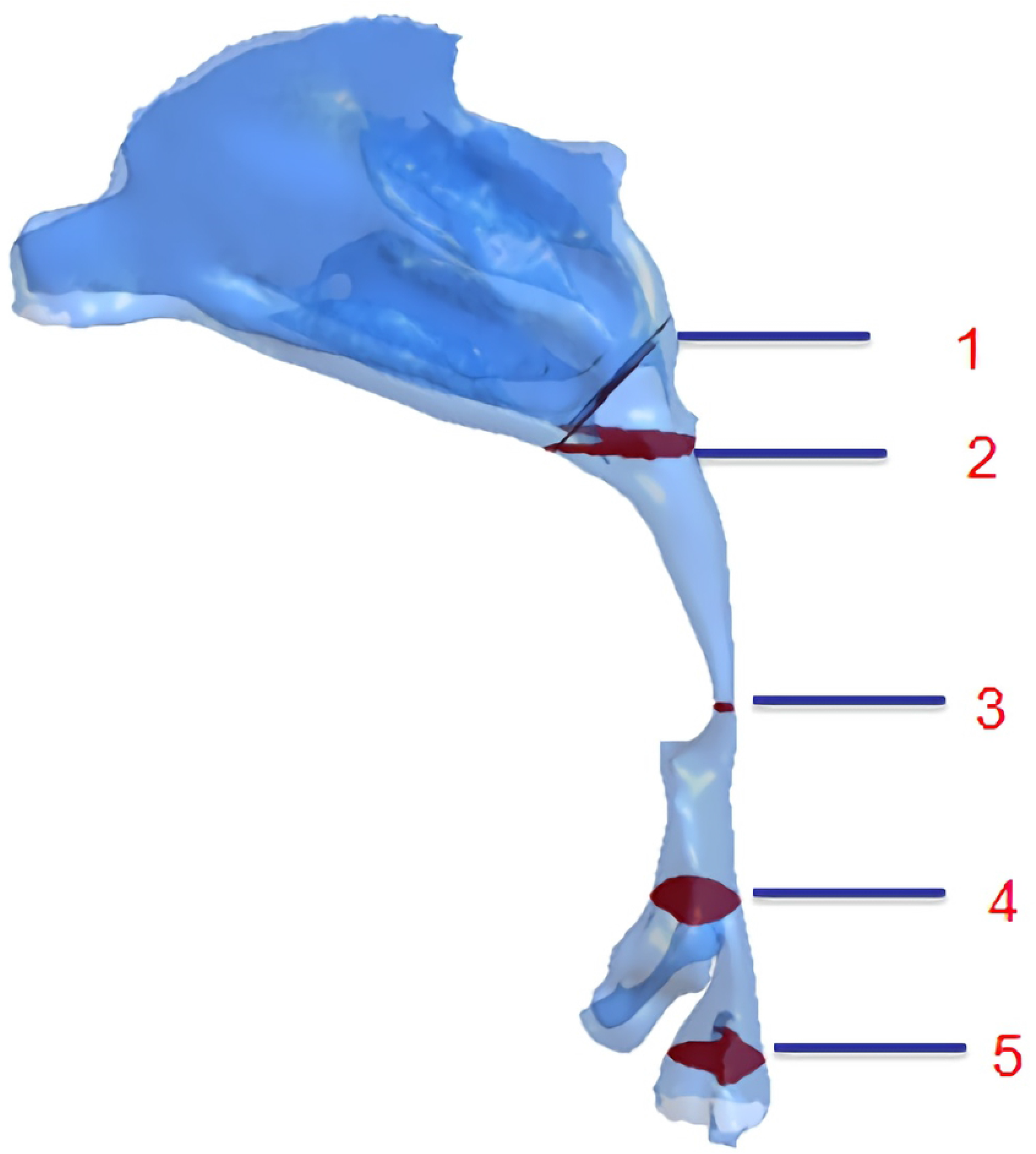
Selected sections of the CFD model.

### Statistical analyses

Statistical analysis was performed using SPSS 17.0 (IBM Corp., Armonk, NY, USA). The paired *t*-test was adopted to compare the changes of the cross-sectional area, volume, velocity, pressure and resistance before and three months after wearing the OAs. The association between AHI change, volume change and resistance change were analyzed using the coefficient of product moment correlation (Pearson’s correlation coefficient). For all analyses, a probability value of *P* <0.05 was considered to indicate differences that are statistically significant.

## Results

### Morphological changes in the upper airways after use of the OAs

As shown in Tables 1 and 2, the volume of the nasopharynx region of patients with and without OAs use showed no significant difference (P>0.05), while that of the palatopharynx, glossopharynx and hypopharynx increased significantly(P<0.05). Prominent changes occurred mainly in the palatopharynx (increased by 39.54%) and glossopharynx (increased by 44.48%). The narrowest region of the upper airway was located at section 3 (the lower bound of the palatopharynx). Notably, its cross-sectional area increased significantly by 129.06% with the OAs(P<0.05).

**Table 1.**
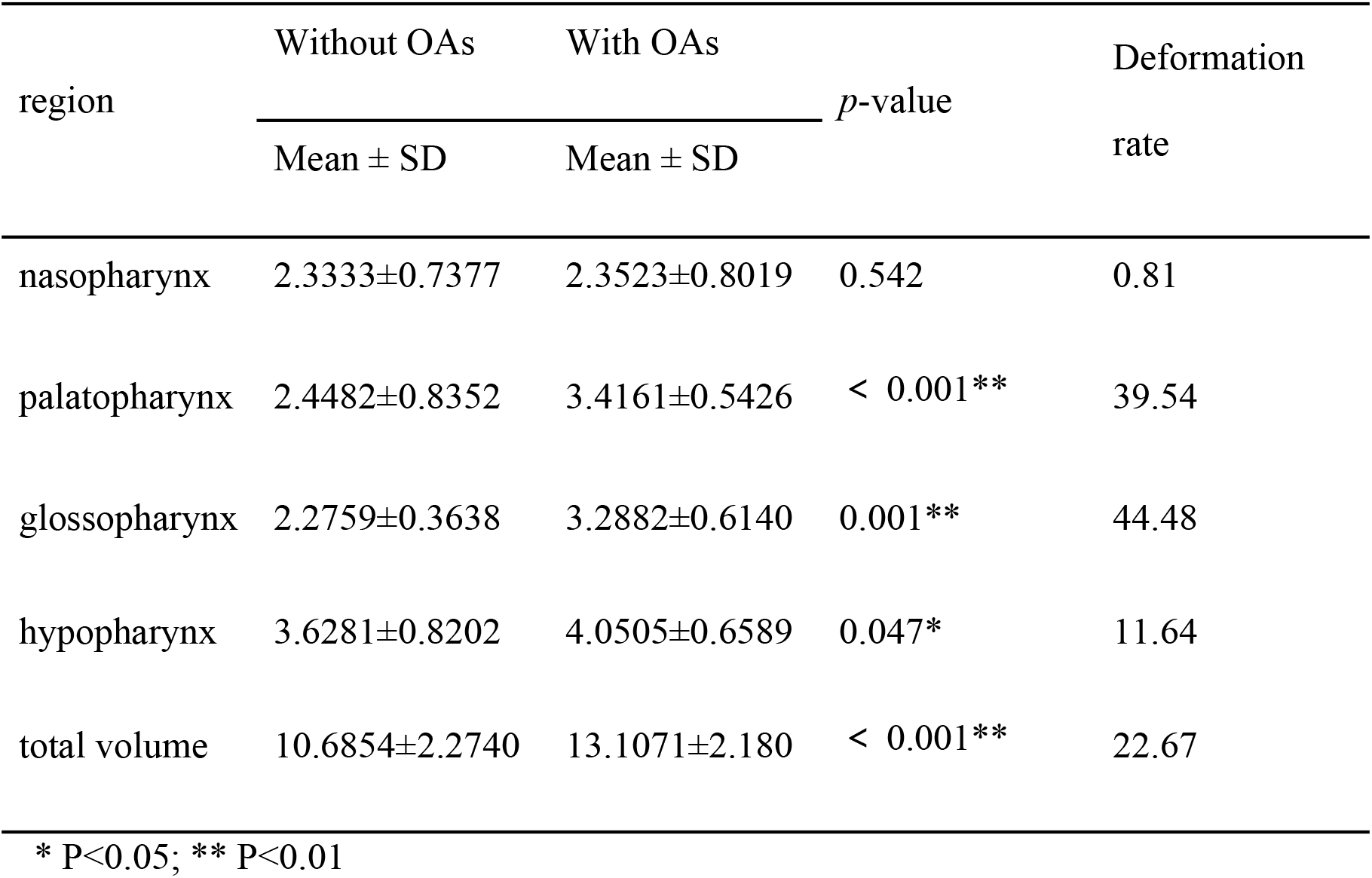
Volumetric changes in the airways with and without the OAs (cm^3^).

**Table 2.** Changes in the cross-sectional area of selected upper airway sections from patients with and without OAs (cm^2^).

| Section | Without OAs | With OAs | p-value | Deformation rate |
| --- | --- | --- | --- | --- |
|  | Mean ± SD | Mean ± SD |  |  |
| A | 3.1441±0.7028 | 3.1643±0.6017 | 0.283 | 0.64 |
| B | 2.1608±0.9456 | 2.3389±0.5536 | 0.058 | 8.24 |
| C | 0.2150±0.1032 | 0.4925±0.1018 | 0.001** | 129.06 |
| D | 1.6430±0.6109 | 2.5817±0.5691 | 0.018* | 57.13 |
| E | 2.3573±0.4370 | 2.6032±0.4092 | 0.035* | 10.43 |

\* P<0.05; \*\* P<0.01

### Airflow velocity changes in the upper airways

The change tendencies of airflow velocity in the UAs of 8 patients were similar to those illustrated in Fig 4. Without the OA, the airflow velocity was turbulent, while it became smooth and steady after using OAs. The maximum airflow velocity in the narrowest part of the pharynx, i.e. the palatopharyngeal region, was determined both pre- and post-treatment. As can be seen in Table 3, there was no difference in the airflow velocity in the nasopharynx and epiglottis pre- and post-treatment (P>0.05), while that of palatopharynx and glossopharynx decreased significantly(P<0.05), from 11.55 m/s to 8.81 m/s post-treatment, representing a decline of 23.7%. The difference was especially significant in the lower bound of palatopharynx (P<0.01).

**Fig 4.**
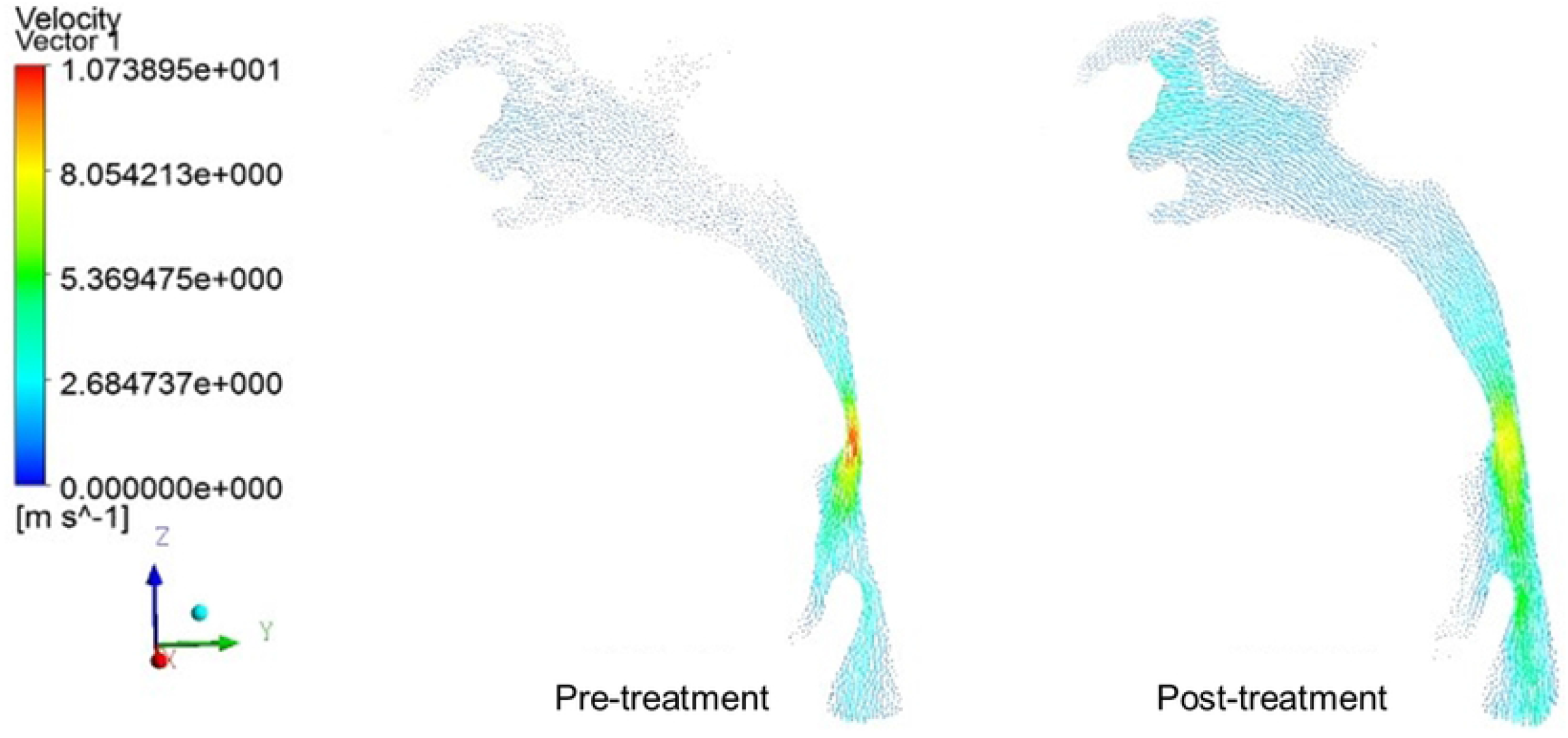
Velocity vectors in the mid–sagittal plane of a patient’s upper airways.

**Table 3.** Airflow velocity changes in different selected sections of the upper airways form patients with and without the OAs (m/s).

| section | Without OAs | With OAs | p-value | Deformation rate |
| --- | --- | --- | --- | --- |
| | Mean ± SD | Mean ± SD | | |
| A | 0.84±0.31 | 0.78±0.24 | 0.183 | -7.14 |
| B | 1.24±0.43 | 1.03±0.55 | 0.048* | -16.94 |
| C | 11.55± 2.18 | 8.81±2.47 | 0.007** | -23.72 |
| D | 4.86±1.17 | 4.34±1.53 | 0.036* | -10.70 |
| E | 2.75±0.74 | 2.60±0.40 | 0.095 | -5.45 |

### Pressure changes in the upper airways of patients before and after using OAs

The pressure change trends in the UA of 8 subjects were similar to what was outlined in Fig 5. The pressure distribution in the UA became steady with OA. Both the initial and final minimum pressure was detected at the lower boundary of the palatopharynx, a narrow region in the UA. Moreover, the maximum flow rate was observed at this point, conforming with Bernoulli’s equation. As shown in Table 4, after wearing the OAs, no statistically significant difference of pressure was observed in the nasopharynx and epiglottis (P<0.05), while pressure in the palatopharynx and glossopharynx increased remarkably(P<0.05), especially at the lower boundary of the palatopharynx(P<0.01), rising from 101240.25 to 101264.13Pa. The pressure in the inlet of the nasal cavity was defined as the baseline (the practical atmospheric pressure is 101325Pa), the pressure at the lower boundary of the palatopharynx without AOs was calculated as −84.75Pa, and it transformed into −60.87Pa with the AOs, thus denoting a pressure drop of 28.2% in the narrowest part of the UAs.

**Fig 5.**
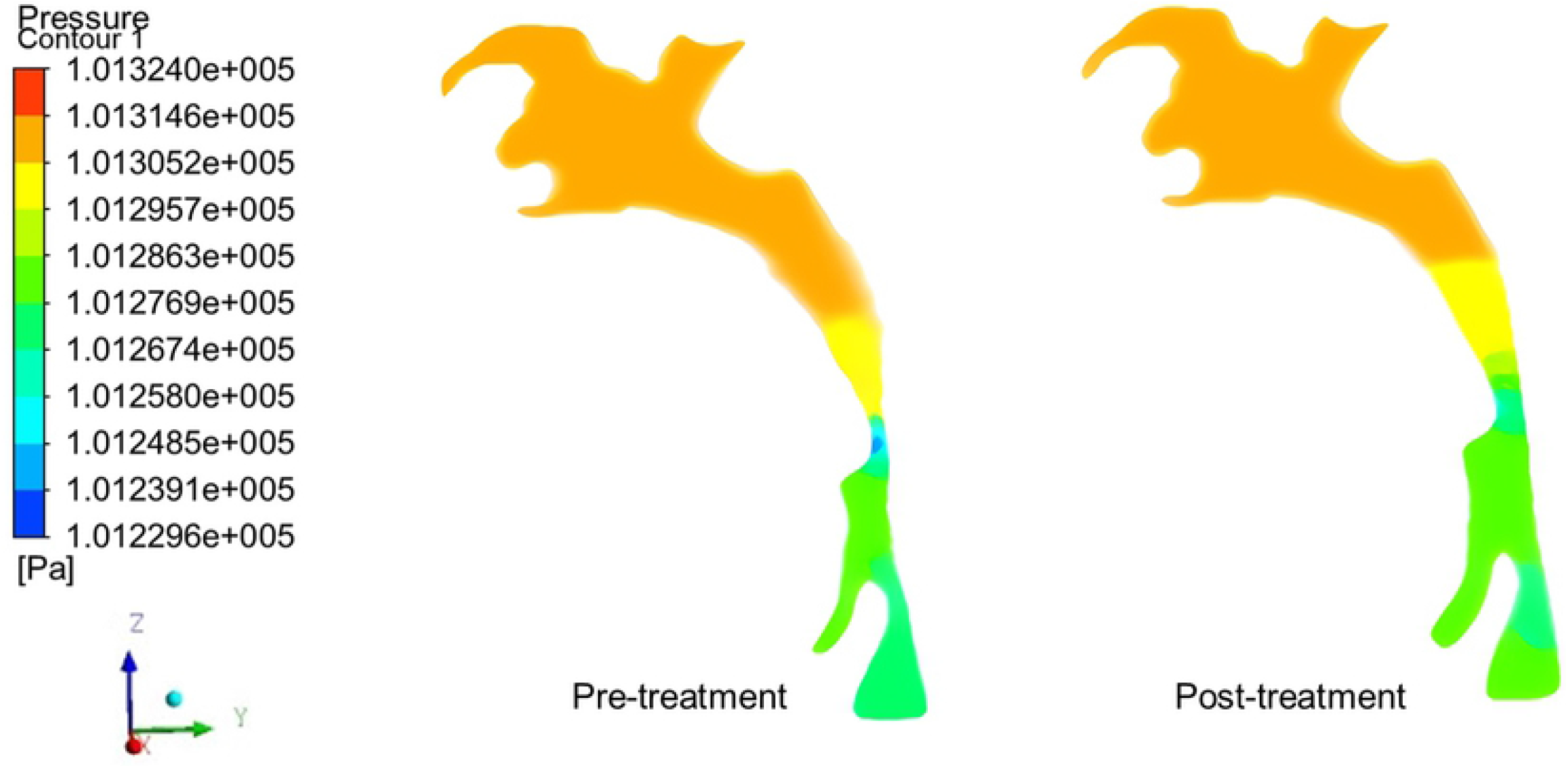
Pressure contours in the mid–sagittal plane of a patient’s upper airways.

**Table 4.** Pressure changes in different sections of the upper airways from patients with and without OAs(Pa).

| Section | Without OAs | With OAs | p-value |
| --- | --- | --- | --- |
| | Mean ± SD | Mean ± SD | |
| A | 101303.61±18.37 | 101305.24±16.79 | 0.093 |
| B | 101308.52±8.46 | 101313.48±6.37 | 0.065 |
| C | 101240.25±16.24 | 101264.13±23.10 | 0.005** |
| D | 101263.49±12.12 | 101272.34±13.47 | 0.039* |
| E | 101255.58±20.66 | 101261.73±18.36 | 0.063 |

### Changes of resistance in the upper airways

Airflow resistance denotes the specific pressure needed to push a defined volume of gas over a designated distance in a defined time. It can be described using the formula: R=∆P/Q, where ∆P indicates the pressure drop, Q indicate the gas flow, which had a set value of 0.2L/s. As shown in Table 5, with the effect of the OAs, the pharyngeal resistance decreased significantly(P<0.05), from 290.63Pas/L to 186.25Pas/L, a decline of 35.9%.

**Table 5.** Changes of resistance in the pharynx of patients with and without OAs (Pas/L).

| Patient | Without OAs |  | With OAs |  |
| --- | --- | --- | --- | --- |
| | $\Delta P$ | R | $\Delta P$ | R |
| 1 | 50 | 250 | 29 | 145 |
| 2 | 42 | 210 | 35 | 175 |
| 3 | 85 | 425 | 32 | 160 |
| 4 | 58 | 290 | 56 | 280 |
| 5 | 82 | 410 | 53 | 265 |
| 6 | 36 | 180 | 33 | 165 |
| 7 | 43 | 215 | 31 | 155 |
| 8 | 69 | 345 | 29 | 145 |
| Mean ± SD | - | 290.63±93.63 | - | 186.25±54.30 |

Note: $t=3.182$ $p=0.015$

### Correlations among AHI, pharyngeal volume and resistance changes in the pharynx with and without OAs

The detailed data on AHI, pharyngeal volume and pharyngeal resistance changes which was collected before and after the 8 patients used OAs is shown in Table 6. Fig 6 shows that there was a negative correlation between pharyngeal resistance changes and pharyngeal volume changes (r= −0.786 p=0.0218, Fig 6-a) as well as pharyngeal volume changes and AHI changes (r= −0.81 p=0.0158, Fig 6-b), while there was a positive correlation between pharyngeal resistance changes and AHI changes (r=0.976 p=0.0008, Fig 6-c).

**Fig 6.**
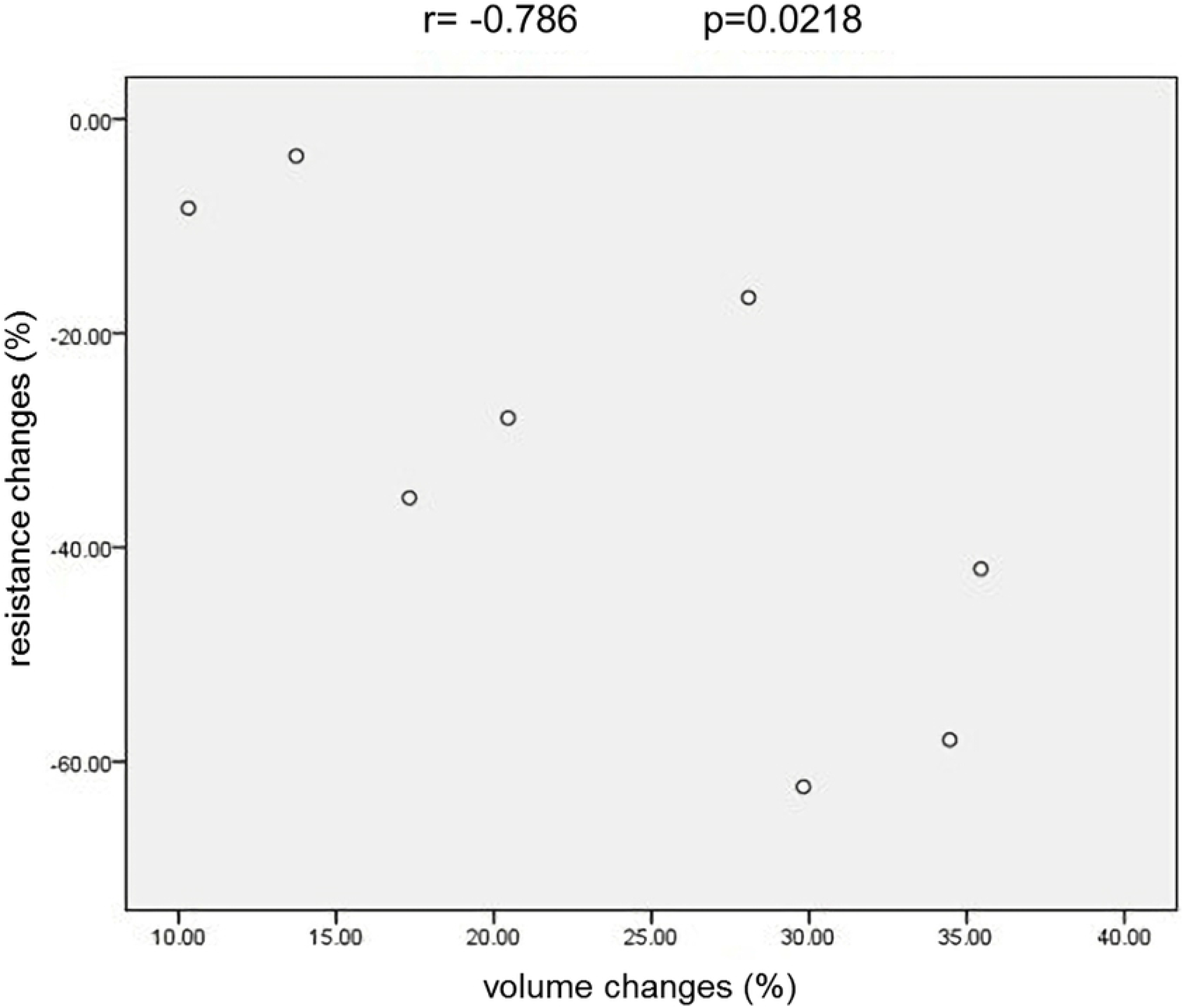
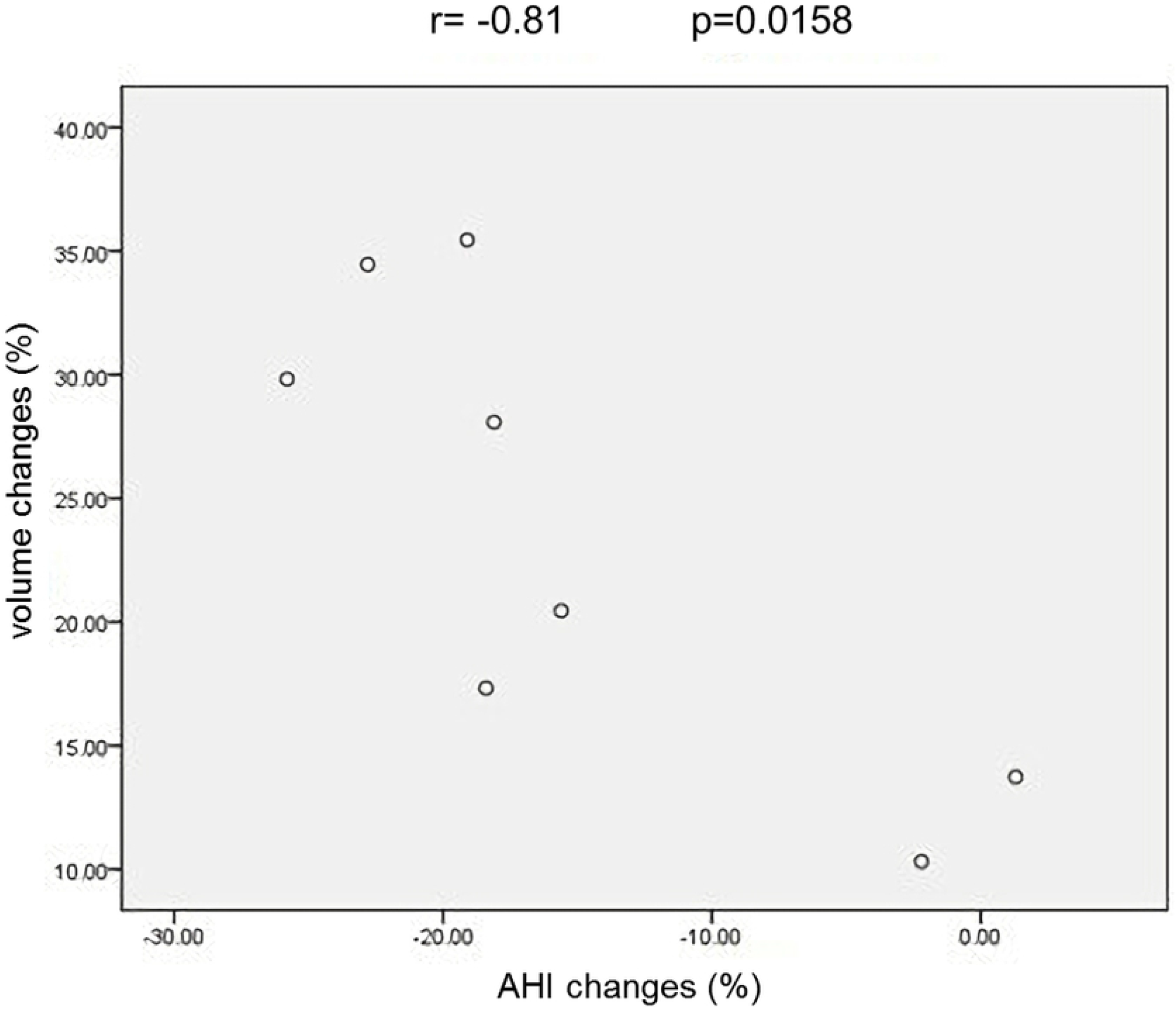
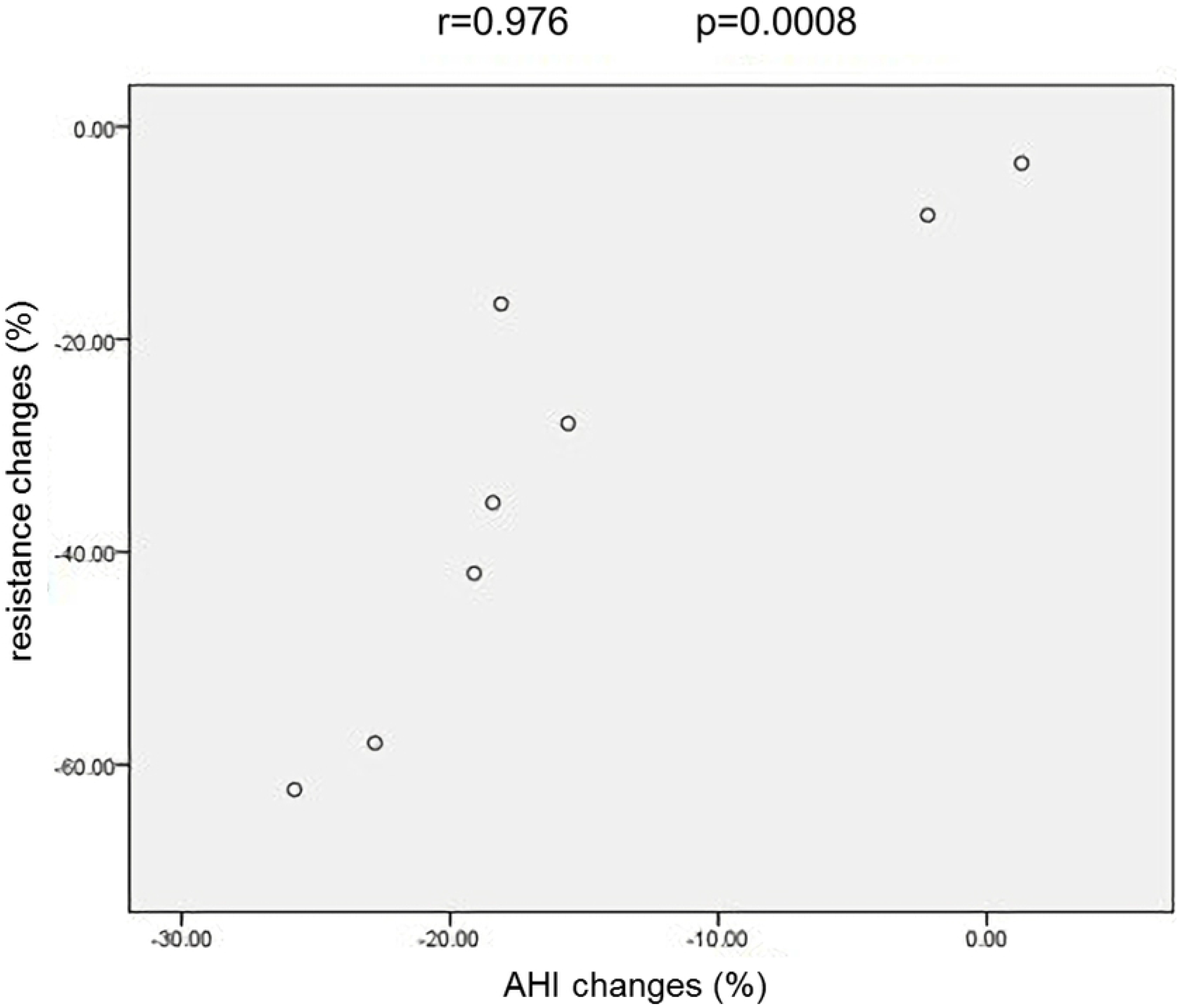
The correlation analysis between the pharyngeal volume changes and pharynx resistance changes(a); between AHI changes and pharyngeal volume changes (b); between AHI changes and pharyngeal resistance changes(c).

**Table 6.** The changes of AHI, pharyngeal volume and pharyngeal resistance in OSAHS Patients with and without OAs (cm^2^).

| patient | Without<br>OAs | With OAs | Deformation rate of<br>pharyngeal<br>volume | Deformation rate of<br>pharyngeal resistance |
| --- | --- | --- | --- | --- |
|  | AHI | AHI | with OAs (%) | with OAs (%) |
| 1 | 21.3 | 2.2 | 35.45 | -42.00 |
| 2 | 23.2 | 5.1 | 28.08 | -16.67 |
| 3 | 27.6 | 1.8 | 29.82 | -62.35 |
| 4 | 16.1 | 16.0 | 13.73 | -3.45 |
| 5 | 21.8 | 3.4 | 17.32 | -35.37 |
| 6 | 14.3 | 12.1 | 10.31 | -8.33 |
| 7 | 17.5 | 1.9 | 20.45 | -27.9 |
| 8 | 24.2 | 1.4 | 34.46 | -57.97 |

## Discussion

### airflow dynamics of the upper airways and OA

The nasopharyngeal airways almost remained unchanged after using OAs, while the cross-section of the glossopharyngeal and palatopharyngeal airways significantly increased, and that of the hypopharyngeal airway increased slightly. These results demonstrate that OAs mainly act on the glossopharyngeal and palatopharyngeal airways. In the narrow region behind the soft palate, the CSA increased obviously by 129.06% (P<0.05). Airway enlargement decreased the possibility of a collapse and obstruction of the UAs during the respiratory process, thus providing better flow conditions. This was confirmed by the analysis of velocity and pressure.

After wearing the appliance, the respiratory flow rate becomes more balanced, especially in the palatopharynx and glossopharynx, where it decreased from 11.55 m/s to 8.81 m/s, resulting in a reduced impact of the airflow on the pharyngeal cavity wall and thus preventing damage to the airway mucosa. The reduced impact will correspondingly reduce the patient’s snoring which arises from high frequency vibrations of soft tissue under the conditions of violent airflow impact on the pharyngeal cavity wall. In the presence of the OA, negative pressure in the narrow area of the palatopharynx decreased significantly from −84.75Pa to −60.87Pa. Within the entire upper respiratory tract, only the palatopharyngeal region consists of soft tissue without bone support, and thus can collapse under the effect of negative pressure. Reducing the negative pressure diminishes airway compliance, and therefore reduces the collapsibility of airways and improves the patients’ symptoms.

Air resistance in the upper respiratory tract is an important index for the evaluation of the ventilation degree, and the resistance value determines the difficulty of inspiration. High resistance indicates that it is difficult for gas to flow in the upper airways, while the converse indicates that it is easy. Resistance in the pharynx decreased significantly by 35.9% (P<0.05), which eased the air circulation in the UAs, further reducing the probability of airway collapse.

By enlarging the upper airways, the OA altered the dynamic characteristics of airflow in the upper respiratory tract, and bringing the morphology into correspondence with function is the fundamental element for the success of OSAHS therapy. Therefore, when devising an OA, one should consider how to achieve changes in airflow pressure and resistance, not merely changes of morphology.

### AHI, total volume and resistance of the pharynx

AHI, introduced in the 1980s, is widely applied as an important indicator in the diagnosis, classification and efficiency evaluation of OSAHS, both in clinical and academic settings. However, investigators found some deficiencies in using AHI to assess efficiency. A study indicates that, after the use of OA, different patients obtained the same AHI, but they suffered from different degrees of hypoxemia^[21]^. Similar studies have also shown that although patients obtain the same AHI after the use of OA, the probability of cardiovascular disease events was actually different^[22]^. Hence, we aimed to identify auxiliary indicators to inspect OA efficacy. We conducted a correlation analysis among the AHI, total volume and resistance of the pharynx before and after the use of OA. There was a negative relationship between pharyngeal volume changes and AHI changes in patients with OA, as well as a positive relationship between pharyngeal resistance changes and AHI changes. Therefore, we extrapolated that the changes of pharyngeal volume and resistance may be auxiliary indicators to test the effect of OAs on OSAHS. Nevertheless, a multitude of clinical data still needs to be gathered and a thorough correlation analysis between these indicators and clinical outcomes needs to be conducted to verify this deduction.

### limitations of the study

Nevertheless, some deficiencies of this study should also be considered. Firstly, the CT data was obtained with patients in the waking state, while respiratory collapse in OSAHS patients mainly occurs during sleep. Secondly, the study ignores the influence of soft tissues, such as muscle tension, which is vital for the stability of the respiratory tract during sleep, as well as changes and movement of the soft palate, tongue root and other soft tissues related to the respiratory tract may be important factors related to the pathophysiology of OSAHS. While in this study we failed to take the effect of soft tissue into consideration, the most likely site of collapse in the respiratory tract depends on respiratory compliance, a certain degree of muscularization, the so-called “tissue stress” and internal respiratory pressure. Therefore, high-resolution CT data and fluid-structure interaction (FSI) numerical simulation technology can be used to further support the present results. However, FSI technology is still in the early research phase and has been applied only to two-dimensional models^[23]^. To be applied in practical research on the oropharyngeal tract, FSI technology needs to overcome a major problem of data availability for particular material properties of tissues such as tongue and soft palate. In addition, the elastic modulus and Poisson’s ratio of tissues measured in vitro may not have a close correlation with tissue deformability in vivo. Therefore, recording CT images of patients in the sleeping state and attaining reliable soft tissue material properties for use in conjunction with Fluid-structure interaction technology(FSI) could enable the analysis of the impact coefficient of soft tissue surrounding the airways and air in the respiratory tract. The effects of these properties on respiratory stability will be the focus of our future research.

In spite of these limitations, our results show that analyzing the dynamic airflow changes in the upper airways in the presence of OAs by applying CFD technology helps us to recognize the mechanism by which OAs exert their effect in OSAHS treatment and the interactive relationship between structure and function of the upper airways, providing theoretical basis for clinical application of OA in OSAHS treatment.

## Conclusions

In OSAHS that received OA treatment, changes in the morphology and flow dynamics of the upper airways were mainly observed in the palatopharyngeal and glossopharyngeal region. When wearing the OA, the upper airway was enlarged, dynamic airflow characters were changed, and negative pressure as well as resistance in the narrow area of the UAs was diminished, thus making the upper airways more resilient to collapse as well as maintaining their patency.

## References

[1] Malhotra A, White DP. Obstructive sleep apnoea. Lancet. 2002;360(9328):237–245.

[2] Young T, Palta M, Dempsey J, Skatrud J, Weber S, Badr S. The Occurrence of Sleep-Disordered Breathing Among Middle-Aged Adults. New England Journal of Medicine. 1993;328(17):1230–1235.

[3] Bixler EO, Vgontzas AN, Ten-Have T, Tyson K, Kales A. Effects of age on sleep apnea in men: I. Prevalence and severity. American Journal of Respiratory & Critical Care Medicine. 1998;157(1):144–148.

[4] Barbé F, Durán-Cantolla J, Sánchez-de-la-Torre M, Martínez-Alonso M, Carmona C, Barceló A, et al. Effect of continuous positive airway pressure on the incidence of hypertension and cardiovascular events in nonsleepy patients with obstructive sleep apnea: a randomized controlled trial. Journal of the American Medical Association. 2012;307(20):2161–2168.

[5] Marin JM, Carrizo SJ, Vicente E, Agusti AG. Long-term cardiovascular outcomes in men with obstructive sleep apnoea-hypopnoea with or without treatment with continuous positive airway pressure: an observational study. Lancet. 2005 Mar;365(9464):1046–1053.

[6] Peppard PE, Young T, Palta M, Skatrud J. Prospective Study of the Association between Sleep-Disordered Breathing and Hypertension. New England Journal of Medicine. 2000;342(19):1378–1384.

[7] Naegele B, Pepin JL, Levy P, Bonnet C, Pellat J, Feuerstein C. Cognitive executive dysfunction in patients with obstructive sleep apnea syndrome (OSAS) after CPAP treatment. Sleep. 1998;21(4):392–397.

[8] Awad KM, Malhotra A, Barnet JH, Quan SF, Peppard PE. Exercise Is Associated with a Reduced Incidence of Sleep-disordered Breathing. The American Journal of Medicine. 2012;125(5):485–490.

[9] Sullivan CE, Issa FG, Berthon-Jones M, Eves L. Reversal of obstructive sleep apnoea by continuous positive airway pressure applied through the nares. Lancet. 1981;317(8225):862–865.

[10] Cartwright RD, Samelson CF. The effects of a nonsurgical treatment for obstructive sleep apnea. The tongue-retaining device. The Journal of the American Medical Association. 1982;248(6):705–709.

[11] Fujita S, Conway W, Zorick F, Roth T. Surgical Correction of anatomical azbnormalities in obstructive sleep apnea syndrome: Uvulopalatopharyngoplasty. Otolaryngol Head Neck Surg. 1981;89(6):923–934.

[12] Riley RW, Powell NB, Guilleminault C. Inferior sagittal osteotomy of the mandible with hyoid myotomy-suspension: a new procedure for obstructive sleep apnea. Otolaryngol Head Neck Surg. 1986;94(5):589–593.

[13] Engleman HM, Asgari-Jirhandeh N, Mcleod AL, Ramsay CF, Deary IJ, Douglas NJ. Self-Reported Use of CPAP and Benefits of CPAP Therapy: A Patient Survey. Chest. 1996;109(6):1470–1476.

[14] Weaver TE, Kribbs NB, Pack AI, Kline LR, Chugh DK, Maislin G, et al. Night-to-night variability in CPAP use over first three months of treatment. Sleep. 1997;20(4):278–283.

[15] Larsson H, Carlsson-Nordlander B, Svanborg E. Long-time Follow-up after UPPP for Obstructive Sleep Apnea Syndrome: Results of Sleep Apnea Recordings and Subjective Evaluation 6 Months and 2 Years after Surgery. Acta Oto-Laryngologica. 1991;111(3):582–590.

[16] Wilhelmsson B, Tegelberg A, Walker-Engström ML, Ringqvist M, Andersson L, Krekmanov L, et al. A Prospective Randomized Study of a Dental Appliance Compared with Uvulopalatopharyngoplasty in the Treatment of Obstructive Sleep Apnoea. Acta Oto-laryngologica. 1999;119(4):503–509.

[17] Haskell JA, McCrillis J, Haskell BS, Scheetz JP, Scarfe WC, Farman AG. Effects of Mandibular Advancement Device (MAD) on Airway Dimensions Assessed With Cone-Beam Computed Tomography. Seminars in Orthodontics. 2009;15(2):132–158.

[18] Kyung SH, Park YC, Pae EK. Obstructive sleep apnea patients with the oral appliance experience pharyngeal size and shape changes in three dimensions. Angle Orthod. 2005;75(1):15–22.

[19] Sam K, Lam B, Ooi CG, Cooke M, Ip MS. Effect of a non-adjustable oral appliance on upper airway morphology in obstructive sleep apnoea. Respir Med. 2006;100(5):897–902.

[20] Ferretti GR, Bricault I, Coulomb M. Virtual tools for imaging of the thorax. European Respiratory Journal. 2001 Aug;18(2):381–392.

[21] Series F, Marc I, Cormier Y, La Forge J. Utility of Nocturnal Home Oximetry for Case Finding in Patients with Suspected Sleep Apnea Hypopnea Syndrome. Ann Intern Med. 1993;119(6):449–453.

[22] Berman EJ, Dibenedetto RJ, Causey DE, Mims T, Conneff M, Goodman LS, et al. Right Ventricular Hypertrophy Detected by Echocardiography in Patients with Newly Diagnosed Obstructive Sleep Apnea. Chest. 1991;100(2):347–350.

[23] Malhotra A, Huang Y, Fogel RB, Pillar G, Edwards JK, Kikinis R, et al. The male predisposition to pharyngeal collapse: importance of airway length. Am J Respir Crit Care Med. 2002;166(10):1388–1395.

